# Bridging laboratory and field research: method adjustments to manipulate field-derived *Aedes aegypti*

**DOI:** 10.1101/2025.07.10.664070

**Authors:** Yanouk Epelboin, Leonardo Daniel Ortega-Lopez, Emilie Balthazar, Alaïs Cornement, Amandine Guidez, Isabelle Dusfour, Mathilde Gendrin

## Abstract

Reference strains of *Aedes aegypti*, reared over numerous generations under laboratory conditions, are selected to fit laboratory conditions and commonly used in research due to their consistency and ease of handling. However, working exclusively with such strains may not fully reflect the traits of natural populations. While working on such strains is relevant in terms of reproducibility between labs, it is also important to perform some work on field-collected mosquitoes and their progeny to capture more representative biological and behavioural variation. In this study, we used the New Orleans reference strain as a control to adjust and evaluate methods to manipulate the F1 progeny of field-collected mosquitoes from Cayenne (French Guiana). To improve blood feeding rate, we tested the impact of several blood feeding systems for mosquitoes kept in a cage or in individual vials and adjusted starvation time. To monitor fertilization, we assessed if dissection buffer affects burst of spermatheca during dissection, whether mosquitoes were collected alive of shortly after death. The results described here may be helpful for studies on mosquito fitness, particularly on field-derived mosquitoes or on experiments requiring individual level monitoring.

**Author summary:** *Aedes aegypti* is the major vector of dengue virus, causing hundreds of millions of cases per year. To fight against disease transmission, mosquitoes are studied in the laboratory, notably using colonies of mosquitoes kept over decades in the laboratory. These reference strains have been selected to fit laboratory conditions and may thus not fully reflect the traits of natural populations. While working on such strains is relevant in a sake of consistency between labs and ease of handling, it is also important to work on field-collected mosquitoes and their progeny to capture more representative biological and behavioural variation. In this study, we adjusted and evaluated methods to manipulate the F1 progeny of field-collected mosquitoes from Cayenne (French Guiana), using the New Orleans reference strain as a control. To improve blood feeding rate, we tested the impact of several blood feeding systems for mosquitoes kept in a cage or in individual vials and adjusted starvation time. To monitor fertilization, we assessed if dissection buffer affects burst of spermatheca during dissection, whether mosquitoes were collected alive of shortly after death. The results described here may be helpful for studies on mosquito fitness, particularly on field-derived mosquitoes or on experiments requiring individual level monitoring.

## Introduction

*Aedes aegypti* is a major vector of arboviruses that strongly impact global health. This mosquito species is implicated in transmission of dengue, yellow fever and the recently emerged chikungunya and Zika viruses. Among them, dengue outbreaks occur annually in tropical regions causing an approximate burden of 390 million cases per year [1,2]. Between 2010 and 2019, chikungunya and Zika viruses increased in prevalence and caused outbreaks of more than 100,000 clinical cases worldwide [3]. While yellow fever has been less prevalent in recent years, it caused an estimate of 109,000 cases in Africa and South America in 2018, half of them leading to death [4]. While vaccines against these arboviruses are under development [5] or exist (yellow fever) [6], vector control is still essential to prevent their transmission [7].

Studying the ecology and biology of *Ae. aegypti* is essential to understand the key factors that contribute to its success as a vector. Applying the knowledge gained from these studies can enhance vector control strategies, making them more targeted and effective in preventing arboviral transmission. By studying how environmental factors such as temperature, humidity, light exposure, nutrient availability, influence on ecological and biological traits such as mating dynamics, population growth and fecundity, vector control measures can be adjusted [8]. Determining causal effects of such environmental factors on life history traits becomes almost impossible with wild populations in field conditions due to natural variability in environmental variables (e.g. temperature, humidity, light, wind) and among individuals – linked to diet, age, genetics, and body size [9]. Therefore, conducting experiments in controlled laboratory conditions is useful to determine whether biological and ecological factors are directly related to environmental variables and to establish the relative importance of environmental variables on life history traits.

Reference laboratory strains of *Ae. aegypti* have enabled to determine standard phenotypic and genotypic traits of individual mosquitoes, which is essential to understanding their biology and their interactions with pathogens. However, despite the advantages of maintaining laboratory mosquito colonies to determine causal influences of the environment on various traits of *Ae. aegypti*, findings are not always transposable to wild-type populations [10]. This discrepancy might be due to long-term adaptations of laboratory colonies that affect their survival and reproductive fitness [11]. Inbreeding, bottle-necks and genetic drift are some of the underlaying causes of such differences [11,12]. Experimental designs must address this issue to ensure findings are relevant and applicable to natural populations, notably for studies focusing on the influence of genetic variability on life history traits or mosquito microbiota.

While most of the earlier studies on mosquito blood feeding were based on the use of live animal hosts, artificial feeding systems are increasingly used to follow the principles of Replacement, Reduction, and Refinement (the 3Rs) [13]. Various artificial feeding techniques were gradually developed including membrane and capillary feeding, based on natural or engineered biocompatible materials [14–16]. Membrane feeding is commonly used when mass-rearing *Ae. aegypti* [17], generally with glass membrane feeders [18] or Hemotek systems (Discovery Workshops, Accrington, UK) which maintain blood temperature [19]. In recent years, several devices made from common and low-cost lab materials have been used to feed mosquitoes [20–22] and appear to be adapted for laboratory mosquitoes [23]. Various blood sources (human, avian, cattle, cow, etc.), and membrane types (chicken skin, collagen, latex gloves and Parafilm®) have been tested to optimize blood feeding [14,16]. Protocols are however not always transferable from reference laboratory strains to field-collected mosquitoes, so method adjustments are required to improve blood feeding.

In this context, this study aimed to investigate the physiological and behavioral differences between the New Orleans laboratory strain and the first laboratory progeny (F1) from a population of *Ae. aegypti* mosquitoes collected in Cayenne, through three key objectives. First, we compared the efficacy of different blood-feeding methods with various types of blood containers. Second, we evaluated the effects of varying durations of sugar starvation on blood-feeding success. Third, we compared the suitability of two dissection media, PBS and *Aedes* saline solution, in maintaining mosquito tissue integrity. Together, these investigations improve the current understanding of the factors influencing experimental outcomes in *Aedes aegypti* derived from the field under controlled conditions. Our low tech methods can be used in conventional laboratory conditions as well as in field-based laboratories.

## Results

### Effect of the strain of mosquitoes on blood feeding rate

We first quantified the difference in blood feeding rate between mosquitoes of the New Orleans reference strain and the progeny of mosquitoes collected as larvae, referred to here as “Cayenne F1”. We offered blood in a Petri dish covered with a Parafilm® membrane and was placed upside down on the cage so that mosquitoes could bite through this artificial membrane, which was kept warm using a hand warmer. We observed a strong effect of the mosquito strain, with a mean feeding rate of 0.7 for New Orleans and only 0.21 for Cayenne F1 (Figure 1A; *β*=-2.07, Z=-5.11, *p*<0.01; see table A in S1 Text for fitted values). Considering the low blood feeding rate of Cayenne F1, we decided to test whether the type of blood meal offered could influence blood feeding success.

**Figure 1.**
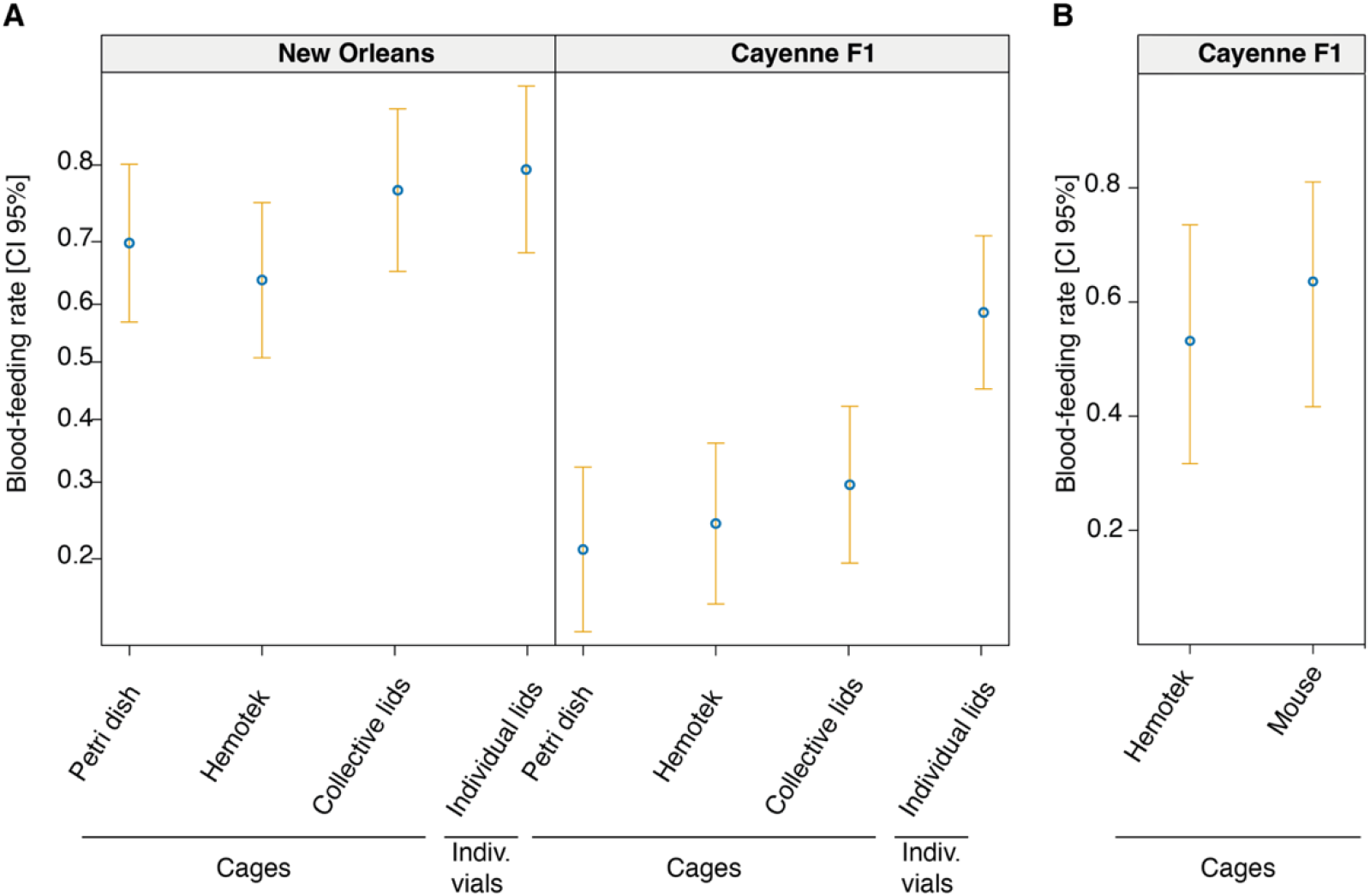
Effect of blood source on the blood-feeding rate of New Orleans strain vs field-derived Ae. aegypti mosquitoes (Cayenne F1) A. Effect of blood-feeding methods on the proportion of blood-fed females after 45min depending on the blood container. All feeding methods deliver human RBCs mixed with human serum; more technical details are provided in the Methods section of the main text. B. Comparison of blood-feeding rates in Cayenne F1 mosquitoes between artificial Hemotek feeding-system containing beef blood and anesthetized mice. Blue dots represent mean predicted values from the final models and yellow bars represent 95% confidence intervals.

### Comparison of blood-feeding techniques

We compared several blood feeding methods: in parallel to the Petri dish method, we offered blood in a Hemotek capsule, in several lids of Eppendorf tubes laid on the cage (‘Collective lids’) or in one lid offered to a mosquito kept in an individual tube (‘Individual lids’, Table 1). This analysis revealed a significant effect of the interaction between mosquito strains and the blood feeding technique on the blood-feeding success (*X*^*2*^_(3)_= 11.26, *p*= 0.01, Figure 1A, Table A in S1 Text), which indicates that the efficiency of blood-feeding methods differed depending on mosquito origin. Specifically, this model predicted a mean feeding rate of 0.73 (95% CI: 0.63-0.81) for the New Orleans strain, and of 0.32 (95% CI: 0.23-0.43) for Cayenne F1. Regarding the blood-feeding method, most Post-Hoc pairwise comparisons did not differ significantly from one another within the New Orleans strain (Table B in S1 Text). However, Post-Hoc pairwise comparisons of blood-feeding methods within the Cayenne F1 strain showed that “individual lids” resulted in the highest blood-feeding rate (0.59, 95% CI: 0.45-0.71;

**Table 1.**
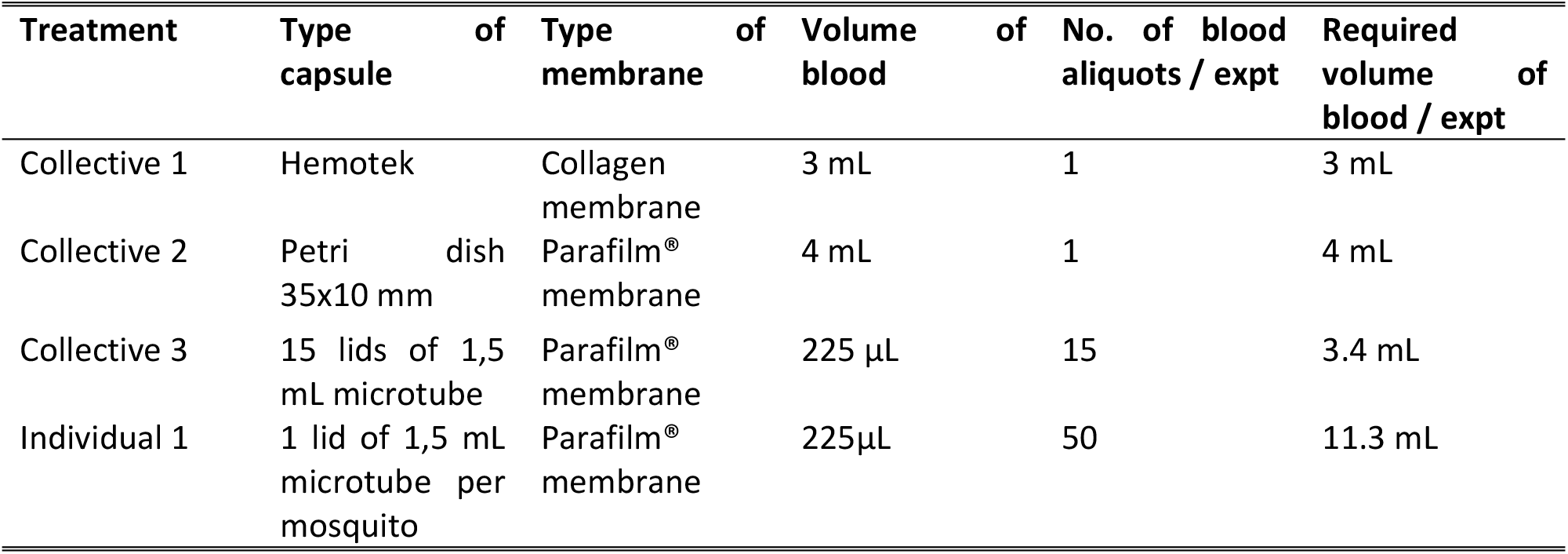
Characteristics of each of the blood-feeding systems used in the experiments.

Figure 1A; *p*<0.001; Table B in S1 Text), compared to the other methods such as collective lids (0.30, 95% CI: 0.20-0.42), Hemotek (0.24, 95% CI: 0.15-0.36), and petri dish (0.21, 95% CI: 0.13-0.32).

We also compared anaesthetized mice and bovine blood in Hemotek as blood-feeding sources. Females of Cayenne F1 *Ae. aegypti* reached a predicted blood-feeding mean rate of 0.64 (95% CI: 0.42-0.81) on mice, slightly higher than their rate of 0.53 (95% CI: 0.32-0.73) on Hemotek devices (*X*^*2*^_(1)_= 4.33, *p*= 0.04, Figure 1B).

### Effect of sugar starvation on blood-feeding rates

Starvation increases blood-feeding rates but implies fitness costs to *Ae. aegypti*. We thus looked for a balance between preventing mortality and maximizing blood-feeding rates. We starved mosquitoes 0 to 48h before offering them blood in Petri dishes. When analyzing mosquito survival, we found that there was no significance from the interaction term between *Ae. aegypti* origin (Cayenne F1 and New Orleans) and any of the four pre-blood-feeding starvation times (*X*^*2*^_(3)_= 1.21, *p*= 0.75). However, we found significant effects from both independent variables separately. Such was the effect of the *Ae. aegypti* strain on blood-feeding rates (*X*^*2*^_(1)_= 4.19, df= 1, p= 0.04; Figure 2A), and the pre-blood-feeding starvation times (*X*^*2*^_(3)_= 111.71, *p*<0.001; Figure 2A). Regarding *Ae. aegypti* strains, New Orleans strain survival rate had a mean predicted value of 0.92 (95% CI: 0.89-0.94) and Cayenne F1 of 0.95 (95% CI: 0.92-0.96) (*β*=0.43, Z=2.03, *p*=0.04). Post-hoc analyses showed that 48 hours of sugar starvation significantly reduced *Ae. aegypti* survival rate (*β*=-2.93, Z=-6.74, *p*<0.001); Table C in S1 Text).

**Figure 2.**
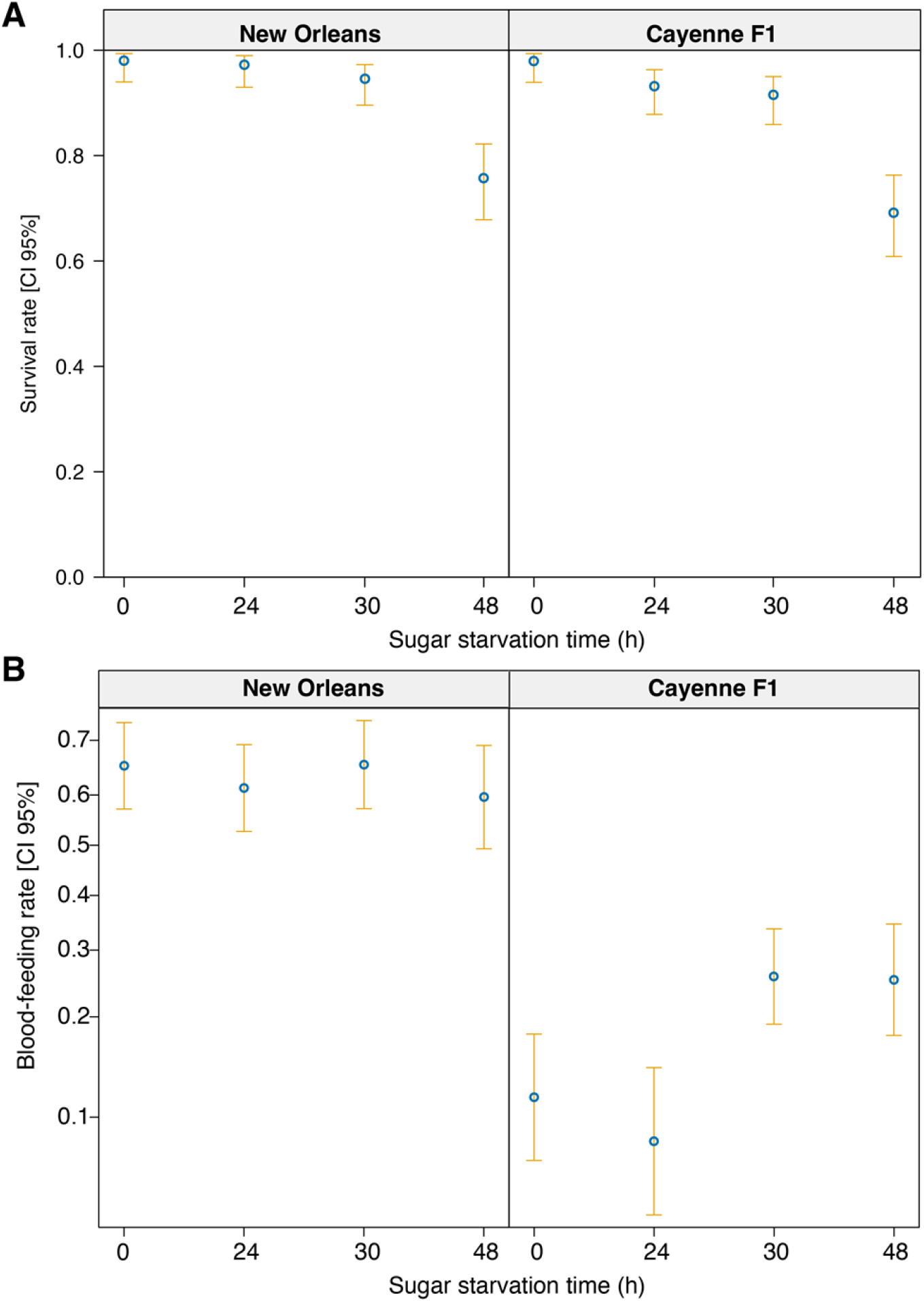
Adjustment of sugar starvation time prior to blood-feeding in New Orleans and Cayenne F1 Ae. aegypti strains. A. Effect of sugar starvation time on mosquito survival at the time of blood-feeding. B. Effect of sugar starvation time on mosquito blood-feeding rate. Blue dots represent the mean predicted values from final models and yellow bars represent 95% confidence intervals.

Considering blood feeding rate, both mosquito strains behaved differently according to starvation time. This effect was seen by observing a significant interaction between *Ae. aegypti* strain and starvation time (*X*^*2*^_(3)_= 14.81, *p*< 0.01, Figure 2B). From Post Hoc analyses, we observed no significant effect between any of the pairwise comparisons of the blood-feeding methods for New Orleans mosquitoes (for all Post Hoc pair-wise comparison results, please see Table D in S1 Text). However, the mean predicted blood-feeding rate of Cayenne F1 was 0.12 at 0 h of sugar starvation (no starvation, 95% CI: 0.07-0.18) and increased to 0.26 at 30 h (95% CI: 0.19-0.34; estimate = -0.98 ± 0.32 SE, Z= - 3.02, *p*= 0.01) and to 0.25 at 48 h (95% CI: 0.18-0.35; estimate = -0.95 ± 0.34 SE, Z= -2.76, *p*= 0.03). Blood-feeding rates at 24 h (0.08, 95% CI: 0.05-0.14) were also significantly lower than at 30 h (estimate = -1.33 ± 0.36 SE, Z= -3.71, *p*< 0.01) and 48 h of sugar starvation (estimate = -1.3 ± 0.38 SE, Z= -3.46, *p*< 0.01). For the rest of the Post Hoc statistical results, please see Table D in S1 Text.

Taken together, these data suggested that starvation was ineffective to increase blood-feeding rates in New Orleans *Ae. aegypti* strain, while a 30 h-starvation was beneficial to increase blood-feeding rates in Cayenne F1 *Ae. aegypti* strain while having little effect on their mortality.

### Effect of dissection conditions on spermatheca integrity

Mosquito *Ae. aegypti* females typically mate once and then carry spermatozoa in their spermatheca. When rearing mosquitoes, it can also be useful to monitor the presence of sperm cells in dissected spermatheca as fertilization status may be associated with behavioural and physiological traits. However, spermatheca sometimes burst during dissection, so we assessed whether the buffer solution affected their preservation. We tested the association between mosquito strain (New Orleans vs Cayenne F1) and the buffer solution used during dissection. Two solutions were compared: phosphate-buffered saline -PBS- and “*Aedes* saline”, which was previously formulated to provide optimal osmotic pressure for mosquito dissections [24].

Assessing fertilization is typically interesting when a colony is struggling, and it is necessary to determine why females are not laying eggs. In such cases, it can be useful to also sample females shortly after death, as live females are sometimes too valuable to be collected. Mosquitoes that fall on the bottom of the cage dry out quickly, but those that drown in water can often be dissected several hours later. Therefore, we also tested whether mosquitoes that had drowned during the night prior to dissection could still be dissected successfully.

We observed that spermatheca integrity was significantly associated with an interaction between dissection buffer and mosquito strain (*X*^*2*^_(1)_= 1.89, *p*= 0.03) as well as with an interaction between dissection buffer and mosquito live/drown status (*X*^*2*^_(1)_= 4.78, *p*<0.001). When analysing mosquitoes as different subsets of the same strain and status, we found that the *Aedes* saline buffer was significantly better than PBS to preserve spermatheca integrity of drown Cayenne F1 mosquitoes (*X*^*2*^_(1)_= 10.02, *p*< 0.01; Figure 3). In contrast, no statistical significance was found for drown mosquitoes of New Orleans strain on spermatheca integrity (*X*^*2*^= 0.45, df= 1, p= 0.33; Figure 3). For mosquitoes sampled alive, both buffers were efficient to preserve spermatheca, as most mosquitoes had 3/3 intact spermatheca (*X*^*2*^_(1)_= 0.1125, *p*= 0.32; Figure 3A).

**Figure 3.**
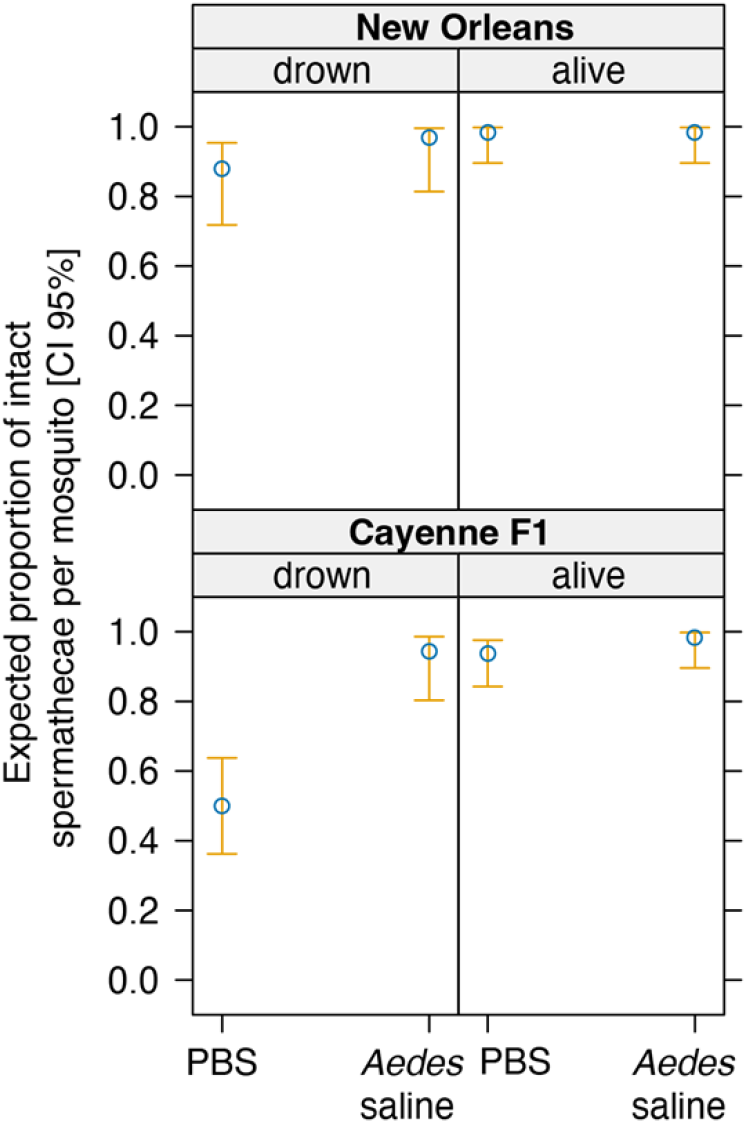
Establishment of experimental conditions for spermatheca dissection. Proportion of intact spermatheca per dissected female from the New Orleans colony (upper panel) or from the progeny of mosquitoes collected in Cayenne (lower panel), depending on the dissection buffer and on the mosquito status upon sampling (drown in the water overnight or live). Blue dots represent mean predicted values from final models and yellow bars represent 95% confidence intervals.

## Discussion

Blood-feeding success of mosquitoes is a fundamental aspect of their fitness and may notably be intended for vector competence studies or to increase the yield of mass rearing. Our findings reveal differences between the New Orleans laboratory strain and the Cayenne F1 field-derived strain of *Ae. aegypti* mosquitoes that have important implications for experimental design, particularly when aiming to maximize blood-feeding efficiency. Laboratory environments generally exert lower selective pressure than field, due to abundant larval diet and, to a lesser extent, controlled temperature and humidity [10]. Nevertheless, *Ae. aegypti* from the field may adapt more efficiently than other insect species from the very first laboratory generation, due to its anthropophilic behaviour and its tendency to breed in artificial containers.

In this study, we presented protocol adjustments aimed at increasing blood-feeding rates in the first progeny of field-collected *Ae. aegypti* mosquitoes. Our results suggest that, regardless of the strain used, a good balance between mosquito survival and blood feeding rate is obtain after a 30-hour starvation using microtube lids dispersed on the grid. We believe these adjustments offer a significant advantage, as this progeny is the closest relative to field mosquitoes. Firstly, we confirmed that the F1 of our local population can feed without live animals by using an artificial blood-feeding device. Feeding on mice was still slightly more efficient than on Hemotek, yet the difference was not strong enough to justify the use of animals, considering ethical reasons and logistical drawbacks. As described in other studies, simple systems can be used with affordable materials, *e*.*g*., cardboard cups, Parafilm-M membrane, and simple heating systems as alternatives to the commercially available Hemotek system [20,22,25,26]. We monitored a similar blood-feeding rate with the use of Petri dish covered with a hand warmer as with Hemotek capsules. Such self-made systems can be particularly useful for several applications, including routine experiments in laboratories, resource-limited settings, or field experiments. Interestingly, lids of microcentrifuge tubes were even more efficient than larger containers to support mosquito feeding. Individualized feeding specifically achieved the highest blood-feeding rate in the Cayenne F1 strain. This could be due to mosquito confinement reducing distance to blood source. We observed a higher success with 15 microtube lids than with a Petri dish while both had a blood surface area of approximately 11 cm^2^. We hypothesize that the dispersion of the 15-lid blood on the cage grid explain this greater success, as it covered overall a greater area. While individualized feeding on microtube lids, whether individualized or not, may not be practical for colony-wide applications, it offers an effective and low-cost approach for experiments requiring precise mosquito development or individual life history traits studies. Additionally, microtube lids limit blood usage compared to other methods, making them particularly useful for resource-limited contexts.

Our data suggest that sugar starvation is unnecessary for the New Orleans laboratory strain but that 30 h of sugar starvation improves blood-feeding rates in the Cayenne field strain (F1). The New Orleans strain may have undergone adaptation to laboratory conditions, enabling more efficient feeding through artificial membranes without requiring additional stimuli such as starvation. Extending sugar starvation time beyond this period did not increase feeding rates further and significantly increased mortality in both strains. Previous studies have shown that sugar starving of 6 to 24 h improves blood-feeding rates in *Aedes* or *Anopheles* [25,27–29]. Conversely, other traits such as host-seeking behaviour, appear unaffected by sugar deprivation in female *Ae. aegypti* mosquitoes [30].

Although a 30-h sugar starvation period improved blood-feeding rates in Cayenne F1 strain, we acknowledge the possibility that this result may be influenced by circadian rhythms of the mosquito, Blood meals were offered at different times for the 30-hour group (3:00 p.m.) compared to the other groups (9:00 a.m.). While activity patterns of *Ae. aegypti* in the field have been described to be bimodal, with peak activity occurring between 7:00 a.m. and 8:00 a.m. and between 7:00 p.m. and 8:00 p.m. [31–34], this species is overall diurnal and can bite throughout the day [35,36]. Both tested times (9:00 a.m. and 3:00 p.m.) fall outside peak activity periods, and further research would be needed to explore whether circadian rhythms contributed to the observed feeding rate differences [38], but the aim of this study was to propose experimental set ups that are convenient in a daily laboratory routine.

The use of adenosine triphosphate (ATP) has been shown to encourage blood-feeding in *Ae. aegypti* [39,40]. ATP cannot be stored at room temperature and is relatively unstable in aqueous solutions, so we did not use this phagostimulant, except for the comparison between Hemotek and live mouse. However, future studies could investigate its effects in combination with other attractants like lactic acid or aldehydes in human sebum. For instance, ketoglutaric acid, or lactic acid are present in animal odours and may be responsible for mosquito attraction [41]. Human sebum composition and long-chain aldehydes such as decanal and undecanal may also be responsible for mosquito preference in host seeking [42]. We did not assess the impact of blood feeding conditions on fecundity and fertility parameters, as studies using similar homemade systems observed no significant differences in egg-laying or hatching rate [16,20,22,26].

Considering conditions to monitor female fertilization, our data indicate that this parameter can still be assessed after death, with females drown overnight. We only observed an effect of the dissecting buffer when Cayenne F1 mosquitoes were sampled as drown, where *Aedes* saline gave better results. These data confirm an advantage of *Aedes* saline for the dissection of sensitive samples [24], yet indicate that PBS is an acceptable general buffer to dissect mosquito tissues. This finding can be useful for monitoring of weak colonies, especially when dissecting specimens post-mortem due to the limited availability of live females in experimental settings.

In conclusion, our study underscores the need to adapt blood-feeding protocols and dissection conditions to the mosquito strain and experimental context. While laboratory-adapted strains like New Orleans perform robustly across feeding systems, field-derived strains such as Cayenne F1 benefit from specific adjustments, including individualized feeding and moderate sugar deprivation. These insights may improve more efficient colony maintenance and experimental reliability, especially in low-resource or field-based laboratories.

## Methods

### Ethics statement

For Figure 1B and some blood meals of New Orleans colony maintenance, anaesthetized mice have been used for mosquito blood feeding. Protocols have been validated by the French Direction générale de la recherche et de l’innovation, ethical board # 089, under the agreement # 973021.

### Mosquito strains

Our reference strain of *Ae. aegypti*, New Orleans, is a well-established laboratory colony routinely maintained in our laboratory (Fx generation). Cayenne F1 mosquitoes are the first progeny of mosquitoes collected as larvae and pupae from 3 to 5 different artificial breeding sites in Cayenne (French Guiana, South America). Larvae were subsequently reared in the laboratory until adult stage, blood fed, mated and were allowed to lay eggs. Multiple larvae collections were carried out to obtain the mosquitoes necessary for the experiments presented here. Both types of mosquitoes are referred to as “strains” throughout this text in a sake of convenience but Cayenne F1 do represent variable local genetic backgrounds.

### Mosquito rearing

Field-collected larvae and pupae from Cayenne were kept in their breeding water and larvae were fed on brewer’s yeast tablets (Gayelord Hauser) until pupal stage. Pupae were removed from breeding trays daily and transferred to plastic containers with tap water in rearing insect cages (Bugdorm®, 30 × 30 × 30 cm) to allow adult emergence. Adult male and female mosquitoes were offered with 10% (w/v) sucrose solution through a soaked cotton which was replaced every 3 days. To obtain eggs, female mosquitoes were blood fed twice a week during 30 minutes on an artificial membrane feeding system Hemotek (6W1, Hemotek Ltd, UK) with heparinized bovine blood provided by the slaughterhouse of Rémire-Montjoly (French Guiana). A plastic container half-filled with dechlorinated tap water and covered on the inner surface with a semi-immerged filter paper was placed inside rearing cages to allow egg-laying. Filter papers were left for 4 days after each blood meal and subsequently removed to dry out in the insectary. Filter papers with eggs were kept dry in zip lock bags until use. Rearing conditions included a daily temperature of 28 ± 3°C, a relative humidity of 80 ± 10% and a natural 12:12 h light:dark cycle.

In order to standardize the age of mosquitoes for each experiment and synchronize hatching, filter papers containing eggs were submerged into about 200 mL of tap water and subsequently hatched inside a vacuum desiccator for 30 minutes. Eggs of Cayenne F1 and New Orleans were hatched separately at the same time. Larvae were then placed in rearing trays and were fed with brewer’s yeast tablets (Gayelord Hauser) until pupal stage. Thus, pupae were collected and transferred into cages for adult emergence. For experiments assessing blood-feeding methods efficiency, adult mosquitoes were sugar fed on cotton soaked in 10% sucrose solution from the first day of emergence until 24 hours before the start of the experiment, or at specific timings for starvation experiments.

### Blood mix preparation

Blood used for experiments corresponds to a 1:1 (v:v) mix of human red blood cells provided by Etablissement Français du Sang (Pointe-à-Pitre, Guadeloupe) with human serum provided by Etablissement Français du Sang (Marseilles, France). Packed red blood cells were washed 3 times with one volume of phosphate buffer saline solution. Supernatant was discarded after a 3900-g centrifugation for 3 minutes.

### Blood-feeding method comparison

Three treatments of group blood feeding and one treatment of individual blood feeding were performed for each of the two mosquito strains (Cayenne F1 and New Orleans). Each experimental trial was performed in triplicate. Three groups of 50 five-day-old females were collected from one rearing cage of each mosquito strain and placed into three separated adult rearing cages for group blood feeding. We used three different blood feeding systems for batch feeding: Hemotek, a Petri dish, and a set of 15 microtube lids (treatment called “collective lids” – meaning collective blood-feeding). Following the volumes described in Table 1, blood was distributed among the feeder devices, which were then covered with the corresponding membranes, placed facing downwards on top of the rearing cages. Female mosquitoes in need of a blood meal could access it by biting through the membrane.

Meanwhile, 50 additional females were also collected from each rearing cage and kept individually for individual blood-feeding. They were allocated into 25 mL plastic vials whose inner surface had previously been scratched with sandpaper to allow mosquitoes to grip, covered with a mosquito net on top. Single microcentrifuge tube lids were filled with 225µL blood, covered with parafilm® membrane and then placed downwards on top of each vial for individual mosquito blood feeding (treatment called “individual lids”-meaning individual blood-feeding).

Except for Hemotek that had its own electronic heating system at 37°C, all other treatments were kept warm with gel hand warmers of sodium acetate that were left on top of the blood-feeding devices after getting activated.

All mosquitoes were offered blood through their corresponding feeding devices for 30 minutes to allow enough time for blood feeding (Figure 4). At the end of the experimental time, all fully engorged females from each treatment were counted.

**Figure 4.**
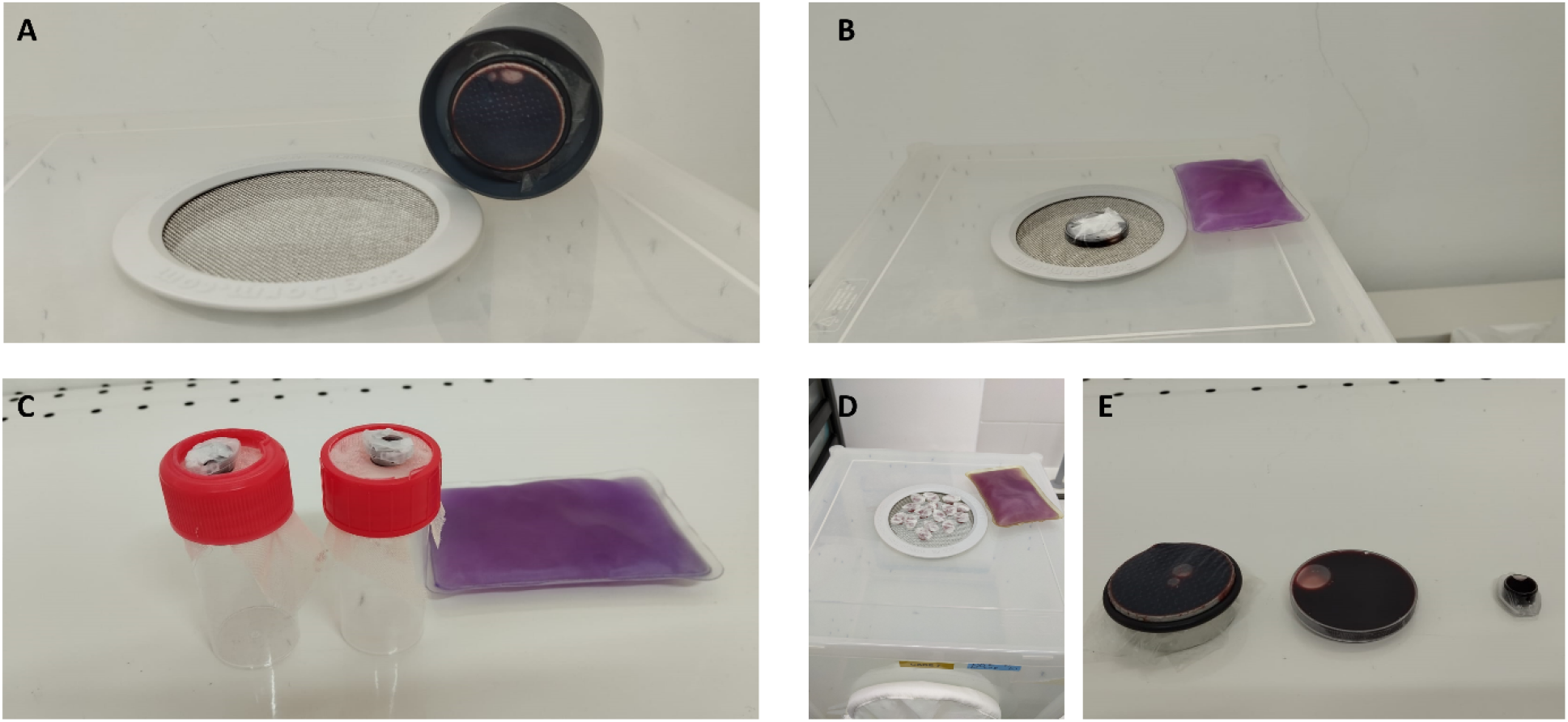
Pictures of the different blood-feeding methods. A: Hemotek system. B: Petri dish of 35×10 mm with a hand warmer. C: Isolated mosquito feeding on an individual 1.5 mL microtube lid with a hand warmer. D: Set of fifteen lids of 1.5 mL microtube with a hand warmer. E: Overview of the three types of blood capsules: Hemotek, Petri dish and lid of 1.5 mL microtube. During blood feeding, the Hemotek membrane lays on the grid in A, and hand warmers lay on top of the blood containers in B-D to maintain temperature.

For blood feeding on mice vs Hemotek, *balb/c* mice (*Mus musculus*) were anaesthetized with a ketamine/xylasine mixture and offered to *Ae. aegypti* females (Cayenne strain) for blood feeding. In parallel, the blood-feeding with Hemotek system was performed in the same condition as described above but instead of human serum, blood was mixed with fetal bovine serum (Sigma-Aldrich) and ATP 3.3% (Sigma-Aldrich). For these experiments, all female mosquitoes were from 7 to 9 days-old post emergence. Experiments were conducted on two separate dates and all mosquitoes were divided into 14 separate cages, with 7 cages for each blood condition. Each cage contained between 80 and 100 females initially and overall, 462 and 493 female *Ae. aegypti* were offered mice and blood in Hemotek respectively. *Ae. aegypti* females were allowed to blood feed for 30 minutes.

### Impact of sugar starvation on blood feeding rates

Four durations of sugar starvation before blood feeding were tested: 0 h (no sugar starvation), 24 h, 30 h, and 48 h. For each treatment, 50 five-day-old adult female mosquitoes from each of the two *Ae. aegypti* strains were selected and placed in separate cages. Depending on the sugar starvation duration, female mosquitoes were either immediately offered a blood meal for 30 minutes or starved for the specified period before being offered the blood meal for the same duration. All females were offered a blood meal using a 35mm-wide and 10 mm-deep Petri dish covered with Parafilm with the corresponding technical details as given in Table 1. Survival was monitored at the start of blood feeding.

### Evaluation of dissection media

Spermathecae of *Ae. aegypti* female were dissected in either Phosphate buffer saline (PBS) at pH=7.4 (tablets from Sigma-Aldrich©) or in “*Aedes* saline” solution at pH= 6.9 (NaCl 9 g.L^-1^, CaCl_2_ 200 mg.L^-1^, KCl 200 mg.L^-1^, NaHCO_3_ 100 mg.L^-1^) [24]. These two solutions were used to determine their effects on tissue integrity and ease of handling during dissections.

New Orleans females were collected from cages where they had spent between 10 and 20 days with males, while field-collected females were not necessarily given time to mate in the laboratory. In both cases, live females were briefly frozen before being dissected under a stereomicroscope with a drop of either of the two buffer solutions. The last abdominal segment was pulled apart with a forceps and opened with a dissecting needle to remove the spermathecae. Integrity of the three spermathecae and presence of spermatozoa were then observed under a phase-contrast microscope with a magnification of 100X.

### Statistical analyses

All statistical analyses were performed using R 4.4.2, and RStudio Pro 2024.04.2. Data were cured and pre-processed in Microsoft Excel, version 16.43 before importing the data into R. All hypotheses were tested using generalized linear mixed models (GLMM), except for survival rate analyses where generalized linear models (GLM) were used. As the response variable for all models was either success (1) or failure (2), for either survival rate or blood-feeding rate, this was fitted to a binomial distribution. Models included the blood-feeding method, the *Aedes* strain, and their interaction as fixed effects, while replicate was included as random effect. These tests were ran using the *lme4* package [43] and figures were all made with the *base* package [44] and modified with Adobe Illustrator 2024. Data were analyzed by building a maximal model to test relevant variables and interactions. Likelihood Ratio Tests (LRT) from a maximal model were applied to assess the contribution of each fixed effect from the model using the *drop1* function from the *stats* package [44]. When categorical variables were significant, Post-Hoc Tukey tests analyses were conducted using *emmeans* and *contrast* functions from the *emmeans* package [45]. More details on statistical methods are provided in S1 Text.

## Supporting information

### S1 text

**Statistical analyses:** details on the different steps for each statistical analysis

**Table B. Estimated marginal means and 95% confidence intervals for the interaction between blood feeding technique (Treatment) and mosquito strain**. *Data was derived from a GLMM fitted with a binomial distribution and a random intercept for Replicate. These values represent predicted blood feeding probabilities and were used to generate the interaction plot in the main manuscript (Figure 1A)*

**Table B. Post hoc pairwise comparisons (Tukey-adjusted) of blood feeding technique (Treatment) within each mosquito strain (Line**). *Data was based on the significant interaction term from the GLMM (maximal model). The table includes estimates, standard errors, z-ratios, and adjusted p-values*.

**Table C. Post hoc pairwise comparisons (Tukey-adjusted) for the effect of sugar-starvation time (Treatment) on mosquito survival**. *Data was based on the GLM (Model 3) after addition of pseudo-observations to address complete separation. The table includes estimates, standard errors, z-ratios, and adjusted p-values*.

**Table D. Post hoc pairwise comparisons (Tukey-adjusted) of blood feeding probability across sugar-starvation treatments (Treatment) within mosquito strains**. *Data was derived from a GLMM with a significant Treatment × Line interaction (Model 4). The table reports pairwise contrasts of estimated marginal means, with corresponding statistics and adjusted p-values*.

